# The Role of Binding Site Specificity in the Disaggregation of A*β*_42_ Fibrils through a Synthetic Paratope

**DOI:** 10.1101/2023.08.11.552930

**Authors:** Shivam Tiwari, Bhubaneswar Mandal, K Anki Reddy

## Abstract

Amyloid-*β* (A*β*) fibrils are the characteristic hallmark of Alzheimer’s disease(AD), and most drug development approaches for AD are focused on preventing and reversing the formation of these fibrillar aggregates. Previous studies show that synthetic antibodies have demonstrated great potential to inhibit the A*β* aggregation and disaggregate the preformed A*β* fibrils. Here, we perform explicit molecular dynamics(MD) simulation to elucidate the molecular mechanism of disaggregation of preformed LS-shaped A*β*_42_ protofibril with a flexible, hairpin-like synthetic paratope (SP) which, in a recent experimental study, has shown promising results. Our simulations demonstrate various potential binding sites for SP on A*β*_42_ protofibril. However, binding of SP at the amyloidogenic core region (KLVFF) shows pronounced structural disruption of A*β*_42_ protofibril. Our results show heavy loss of *β* sheet content, dismantling of K28-A42 salt bridge, and destruction of key contacts in the hydrophobic cores of A*β*_42_ protofibril in the presence of SP. We found the aromatic and hydrophobic residues of A*β*_42_ protofibril participating primarily in the binding with SP. Also, we found that *π* − *π* stacking and hydrophobic interactions are the most dominant mode of interaction between SP and A*β*_42_ protofibril. This work provides a detailed atomistic perspective on the A*β*_42_ protofibril disaggregation mechanism with SP, and the findings can help develop more effective drugs for AD in the future.

## 1 Introduction

Despite decades of extensive research, Alzheimer’s disease (AD) continues to pose a significant challenge in the field of neurodegenerative disorders. The exact mechanism and cause of AD remain elusive. However, one of the most common pathological hallmarks of AD is the presence of amyloid beta (A*β*) plaques and neurofibrillary tangles of Tau proteins(*1*) in the brains of AD patients. Therefore, misfolding and aggregation of A*β* peptide are widely considered one of the most accepted hypotheses for the cause of AD(*1–5*).

A*β* peptide is produced by the cleavage of amyloid precursor protein(APP) by *β* secretase and *γ* secretase(*2*). The splitting of APP by *γ* secretase produces A*β* peptides of varying lengths such as A*β*1-36 to A*β*1-43(*6*), A*β*4-42 and A*β*5-42(*7*), A*β*1-26 and A*β*1-30(*8*). However, the fibrils of species A*β*1-40 and A*β*1-42 are the major constituents of the senile plaques found in AD patients’ brains(*2*). Among the two, A*β*1-42 is less abundant but is higher in toxicity and aggregation propensity than A*β*1-40(*9*). The formation of mature A*β* fibrils occurs through a complex multistep self-assembly process through a nucleation-condensation and polymerization mechanism, which involves multiple intermediate metastable species such as oligomers and protofibrils, which are also toxic in nature(*10–13*). A*β* fibril also exhibits structural polymorphism demonstrated by various studies reporting models having U-shaped, S-shaped, and LS-shaped topology(*14–17*).

The finding “A*β* fibrillation is the primary cause of AD” revolutionized the field of AD research and led all the drug development and therapeutical approaches toward finding a way to inhibit and reverse the A*β* fibrillation process. These approaches involve experimental investigation of many potential inhibitors such as nanoparticles(*18–20*), peptides containing amyloidogenic core region(KLVFF) of A*β* and other short peptides(*21–23*), small molecules(*24–26*) and antibodies(*27, 28*). Furthermore, these experimental studies also inspired many simulation studies(*29–33*) to deepen our understanding of the interaction of the drug and A*β* and the mechanism of inhibition/disaggregation of A*β* fibrils at a molecular level. Lemkul et al. performed an MD simulation to study the molecular interactions involved in the destabilization of A*β* fibrils by a flavonoid called morin. They found that morin can block the attachment of a new peptide by binding at the end of the preformed A*β* fibril. Also, morin diffuses to the core of the A*β* fibril, consequently disrupting crucial hydrophobic interactions(*29*). Viet et al. used MD simulations to study the inhibition of A*β* oligomerization by two breaker peptides, KLVFF and LPFFD. Although both breaker peptides showed inhibitory effects, LPFFD exhibited stronger interference with aggregation and higher binding affinity to A*β*16–22, attributed to favorable hydrophobic interactions(*30*). Agrawal et al. employed MD simulations to study the disruption of U-shaped A*β*40 trimer by 12-crown-4 ether. The study revealed that the 12-crown-4 ether enters the hydrophobic core region and causes the loss of *β* sheet content by interacting strongly with key hydrophobic residues. Also, it destabilizes the Asp23-Lys28 salt bridge by interacting with Lys28(*31*). Zhang et al. used explicit MD simulations to study the interaction between graphene nanosheet and preformed A*β* fibrils. The graphene sheet is found to be interacting strongly with A*β* fibrils leading to structural deformation, particularly for residues having outer side chains. The van der Waals forces were recognized to be the dominant driving force for A*β*-graphene binding, while solvent was found to mediate the interaction between them(*32*). Zhan et al. conducted explicit atomistic simulations to understand the mechanism behind the disruption of A*β*42 protofibril by green tea extracts epigallocatechin-3-gallate(EGCG) and epigallocatechin(ECG). The work concluded that the EGCG exhibited higher disruptive capacity than EGC, owing to the presence of the gallic acid ester group in EGCG. The study sheds light on various atomistic interactions through which EGCG and EGC interact with A*β*42 protofibril such as *π*-*π*, cation-*π*, hydrophobic and hydrogen bonding interaction(*33*).

Recently, Paul et al.(*34*) designed and synthesized a flexible, hairpin-like synthetic paratope(SP) as a potential drug candidate for inhibiting the aggreagation of A*β* and disaggregating the preformed fibrils. A paratope is a part of the antibody which recognizes and binds to the epitope region of the antigen. The SP’s design was inspired by a peptide fragment(LVFFA) of A*β*. The study also tested the efficacy of the newly developed drug(SP) against the inhibition of A*β* aggregation and disaggregation of preformed fibrils and the results were quite promising. Herein, we performed explicit MD simulation to investigate the disassembly mechanism of A*β* fibrils via interaction with SP at a molecular level. In our simulations we observed significant disruption of A*β* fibrils in the presence of SP. Specifically, we analyzed the *β* sheet content of A*β* fibrils and found to have a marked reduction in it due to SP binding. Moreover, the salt bridge between K28 side-chain and terminal residue A42’s COO^−^ group, a crucial stabilizing force for the structural integrity of A*β* fibril, was found to have disrupted in SP’s presence.

We were able to identify locations of various binding sites of SP on A*β* fibril and the participating residues of A*β* and SP. In particular, hydrophobic and aromatic residues were found to be playing the central role in the binding of SP on A*β* fibril. *π* − *π* interactions were found to be the dominant mode of interaction between A*β* fibril and SP.

## 2 Results and Discussion

We have performed six independent 600 ns MD simulations for two sets of systems: 1) A*β*_42_ (2) A*β*_42_ + SP. Figure 1a and 1b shows the atomistic models of isolated A*β*_42_ and A*β*_42_ + SP systems. The A*β*_42_ protofibril model used in this study is a full-length (1-42) LS-shaped A*β* protofibril, which is a nonamer(containing nine chains) in the form of two separate protofibrils, one with five chains(pentamer) and another with four chains(tetramer) as shown in the Figure 1a. Figure 1c shows the key structural features of the LS-shaped A*β*_42_ protofibril model used in the present study. The overall structure of the protofibril is stabilized by three hydrophobic cores (Figure 1c), and the residues forming the stabilizing hydrophobic contacts inside these cores are as follows: (1) core-1: A2, F4, L34, and V36 (2) core-2: L17, E19, and I31 (3) core-3: A30, I32, M35, and V40.

**Figure 1:**
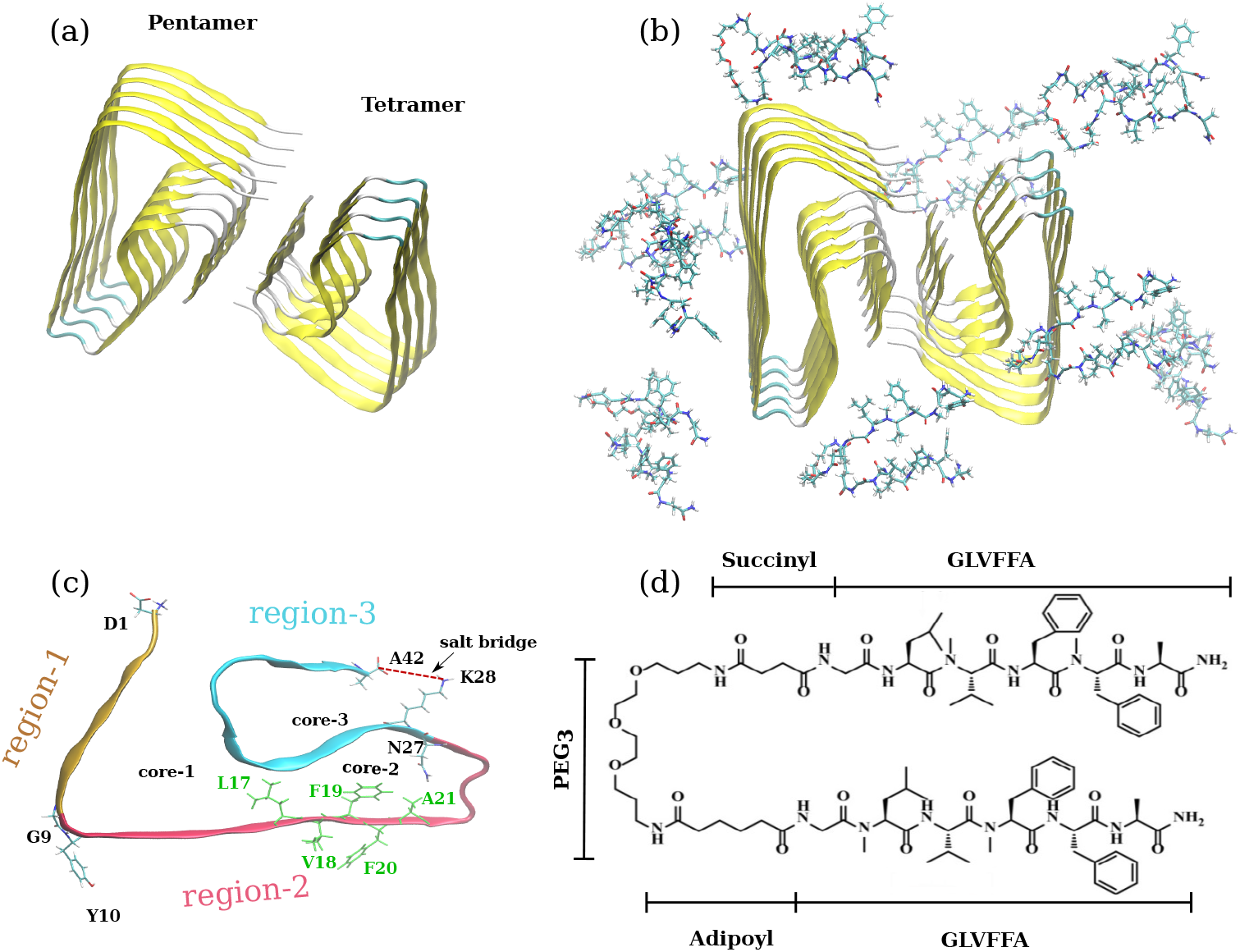
(a) LS-shaped A*β*_42_ nonamer(pentamer + tetramer) protofibril obtained from cryo-EM resolved structure(PDB ID: 5OQV). (b) The initial configurtation of A*β*_42_ + SP system with SP molecules randomly placed around A*β*_42_ nonamer. (c) Structural features of LS-shaped A*β*_42_ protofibril shown on a single chain. The protofibril structure is divided into three regions region-1(D1-G9), region-2(Y10-N27) and region-3(K28-A42) encapsulating the three hydrophobic cores. The residues shown in the green color(LVFFA) belongs to the amyloidogenic region of the A*β*, these residues are also the core ingredients of the ligand(SP) used in the study. The salt bridge between K28 and A42 shown by red dotted line plays a crucial role in the stability of the A*β*_42_ protofibril. (d) Molecular structure of the ligand, a hairpin like synthetic paratope where two strands of the amyloidogenic region(LVFFA) of A*β* are joined by a flexible chain.

The salt bridge between the positively charged 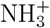 group of K28’s side chain and negatively charged COO^−^(C-terminus) of A42 is also an important interaction for the A*β*_42_ protofibril’s structural stability. Figure 1d shows the chemical structure of the SP molecule. The most crucial part of the SP’s structure is the sequence “LVFFA,” chosen from the central hydrophobic core(CHC) region of A*β*_42_ protofibril(Figure 1c). The reader is referred to the Methods section for a detailed discussion about the ligand design and system preparation.

### SP Disrupts the Structure of A*β*_42_ Protofibril, with Severe Structural Destruction of Tetramer

Root-mean-squared-deviation(RMSD) is a frequently used measure for assessing the structural stability of proteins, and any perturbation to the protein’s structure will reflect in the RMSD. We calculated the time evolution of RMSD for C_*α*_ atoms of A*β*_42_ protofibril for tetramer and pentamer in isolated A*β*_42_ system and A*β*_42_ + SP system with respect to their corresponding initial structure. From now on, we will refer to the isolated A*β*_42_ system as control and A*β*_42_ + SP system as ligand unless otherwise stated. Figure 2a, and 2b compares control and ligand’ C_*α*_ RMSD of A*β*_42_ for tetramer and pentamer, respectively. The RMSD analysis shows that the SP causes profound disruption of A*β*_42_ protofibril’s structure in the case of tetramer(Figure 2a). The RMSD for the ligand system in tetramer shows a sharp increase during the first 50 ns and then converges and fluctuates around 0.2 nm up to 300 ns; during this interval(50 ns - 300 ns), the RMSD does not show any drastic changes and remain in between 0.2-0.25 nm. However, after 300 ns, we observe an acute increase in the RMSD, reaching up to 0.5 nm, indicating a severe disruption of the tetramer’s structure in the presence of SP. Contrarily, the tetramer’s RMSD in the control system increases quickly during the beginning of the trajectory and then converges at around 0.12 nm. The RMSD suggests that the structure of A*β*_42_ tetramer in the control system remains stable throughout the trajectory, except for a slight increase at around 200 ns and 370 ns, the RMSD stays within 0.12-0.19 nm. In the case of pentamer, the RMSD(Figure 2b), for both the control and ligand system, stabilizes at around 0.15 nm after an initial increase. The RMSD analysis for the pentamer reveals that the presence of SP does not have a marked influence on the structure since the RMSD for the ligand system shows no noticeable deviation from that of the control system. Furthermore, a comparison between the RMSD of the tetramer and pentamer in the control system shows that the pentamer’s RMSD stabilizes at 0.15 nm and that of the tetramer at 0.19 nm, implying that the pentamer possesses higher inherent stability than the tetramer.

**Figure 2:**
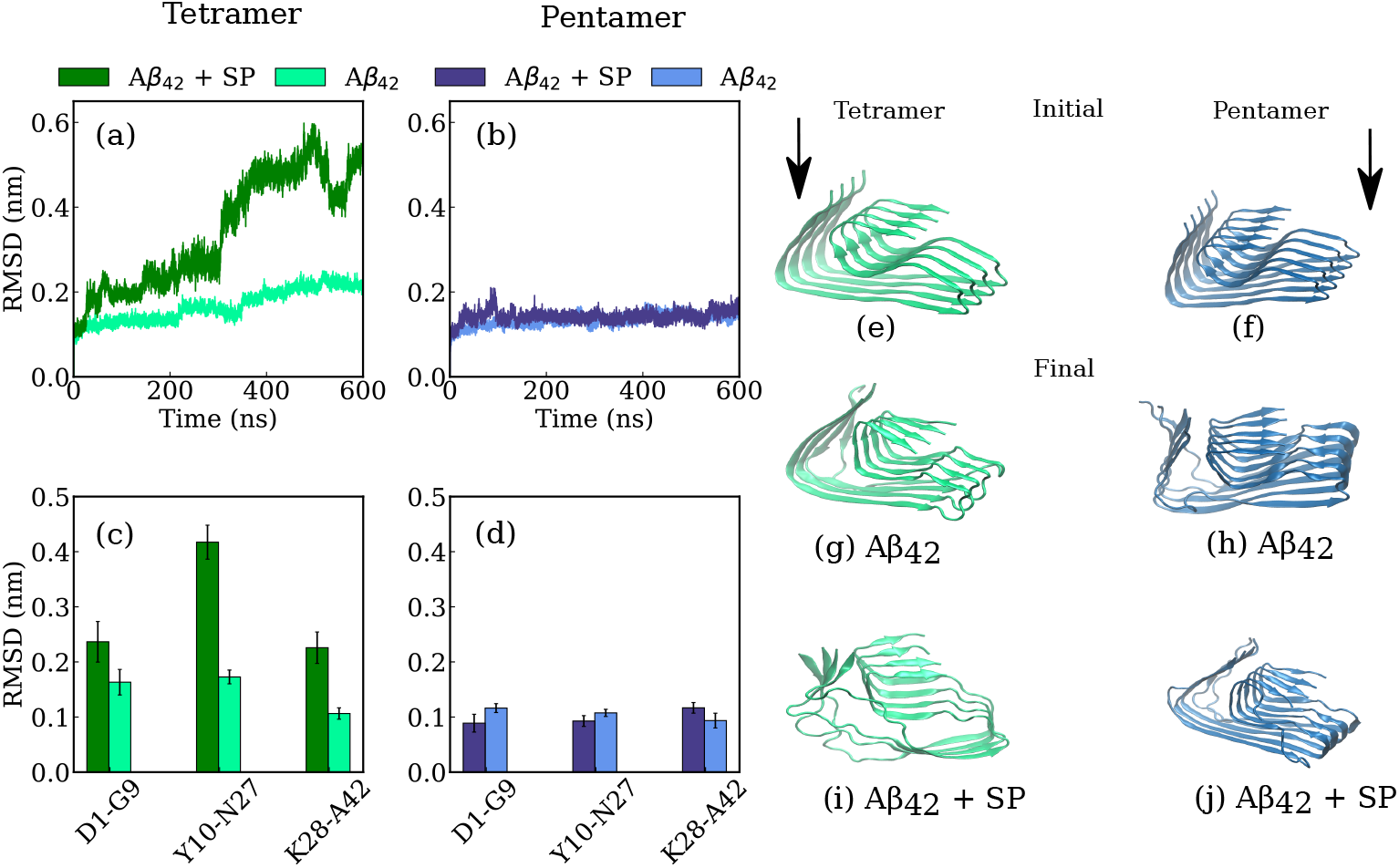
Time evolution of C*α* RMSD of (a) Tetramer in control(light green) and ligand(dark green) system (b) Pentamer in control(light blue) and ligand(dark blue) system. C*α* RMSD values of different regions of A*β*_42_ protofibril averaged over last 200 ns for the (c)Tetramer in control(light green) and ligand(dark green) system (d) Pentamer in control(light blue) and ligand(dark blue) system. Initial structure of A*β*_42_ protofibril of (e) Tetramer and (f) Pentamer. Structures of A*β*_42_ protofibril at the end of the simulation time (g) Tetramer (h) Pentamer in the control system and (i) Tetramer (j) Pentamer in the ligand system.

To further illuminate the effect of SP on the structural stability of A*β*_4_2 protofibril and to explore its most affected region due to the presence of SP, we calculated the average RMSD of different regions in the protofibril’s structure for pentamer and tetramer. The three selected regions are shown in Figure 1c. The selected regions are region-1: D1-G9, region-2: Y10-N27, and region-3: K28-A42. Figure 2c shows the average RMSD for tetramer in the control and ligand system. The RMSD suggests instability in all three regions of tetramer in the presence of SP, indicated by higher RMSD in the ligand system. Interestingly, region-2(Y10-N27) in the tetramer shows the highest RMSD(0.42 nm). Region-2 contains the CHC(LVFFA) of A*β*_4_2, which is also the key ingredient in the SP’s design and is known to be a self-recognition unit. Hence, the high RMSD of region-2 indicates that the SP can recognize the CHC region and bind around it. However, in other sections, we will discuss the specific binding site of SP on A*β*_4_2. The RMSD for D1-G9 and K28-A42 regions for control/ligand was found to be 0.18/0.22 and 0.21/0.10, respectively. RMSD analysis for tetramer shows that SP significantly destabilizes all three regions of the protofibril. Contrarily, pentamer’s RMSD(Figure 1d) shows that SP’s presence has minimal effect on the protofibril’s structure. The RMSD(control/ligand) for region D1-G9(0.11/0.09) and Y10-N27(0.11/0.09) shows a slight decrement in the presence of SP, suggesting that SP is stabilizing the protofibril in the case of pentamer. Conversely, the K28-A42 region in the pentamer shows a little increase in RMSD(0.09/0.11) in the ligand system, indicating a slight perturbation of the structure due to SP. The RMSD analysis reveals that the presence of SP strongly affects the tetramer’s structure, particularly the CHC-containing region-2 is severely affected. However, pentamer do not show any significant structural changes in the presence of SP. A visual analysis of the initial and final conformation of pentamer and tetramer protofibril(Figure 1e-j) also agrees with the results obtained by the RMSD analysis.

### SP Destroys Tetramer’s *β* Sheet Structure

*β* sheets are the characteristic structural features of amyloid beta fibrils. Hence, the disruption of ordered *β* sheets is a crucial indicator of the efficacy of a potential drug aiming to disaggregate the A*β* fibrils. In order to evaluate the impact of SP binding on the *β* sheet structure of A*β*_42_ protofibril, we calculated per-residue *β* sheet probability for tetramer and pentamer in control and ligand systems. Interestingly, the *β*-sheet probability analysis of tetramer(Figure 3) shows a substantial loss of *β*-sheet content, in the presence of SP, around the region-2 of the protofibril. Specifically, residues between Y10 and F19(Y10, E11, V12, H13, H14 and F19) shows drastic reduction in the *β* sheet probablity. These observations for region-2 reveal crucial information about the SP binding and the protofibril disaggregation. The residues observing the highest reduction in the *β*-sheet probability are all sequential(Y10-H14), implying that the disaggregation process primarily occurs around these residues; this can be verified visually from Figure 2e and 2i. It can be observed that the region around core-1 has been disrupted severely, and the *β*-sheets have turned to random coils for all the chains in the tetramer. Moreover, the heavy loss of *β*-sheet content in sequential residues suggests that the SP preferentially attacks region-2 and probably enjoys the strongest binding around these residues(precise details of SP binding will be discussed in another section). However, it can also be observed from Figure 3a that there is an increase in *β*-sheet probability for some residues in the tetramer, such as D1, E3, E22, D23, and G29. In contrast, the *β*-sheet probability for pentamer(Figure 3b) shows an increase in *β*-sheet content for most of the residues in the presence of SP. The residues showing a significant increase in *β*-sheet probability are E22, D23, and V24, and residues between S8-H14. Intriguingly, S8-H14 is the same region that witnessed a drastic reduction in the *β*-sheet content for tetramer in the presence of SP. Another interesting observation here is that the *β*-sheet around the C-terminal region(A30-V40) remains stable and relatively unaffected by the SP’s presence in case of both tetramer and pentamer. Overall, the *β*-sheet probability reveals interesting details about residues and the regions of A*β*_4_2 protofibril, involved in the disaggregation process and about the probable region of SP binding.

**Figure 3:**
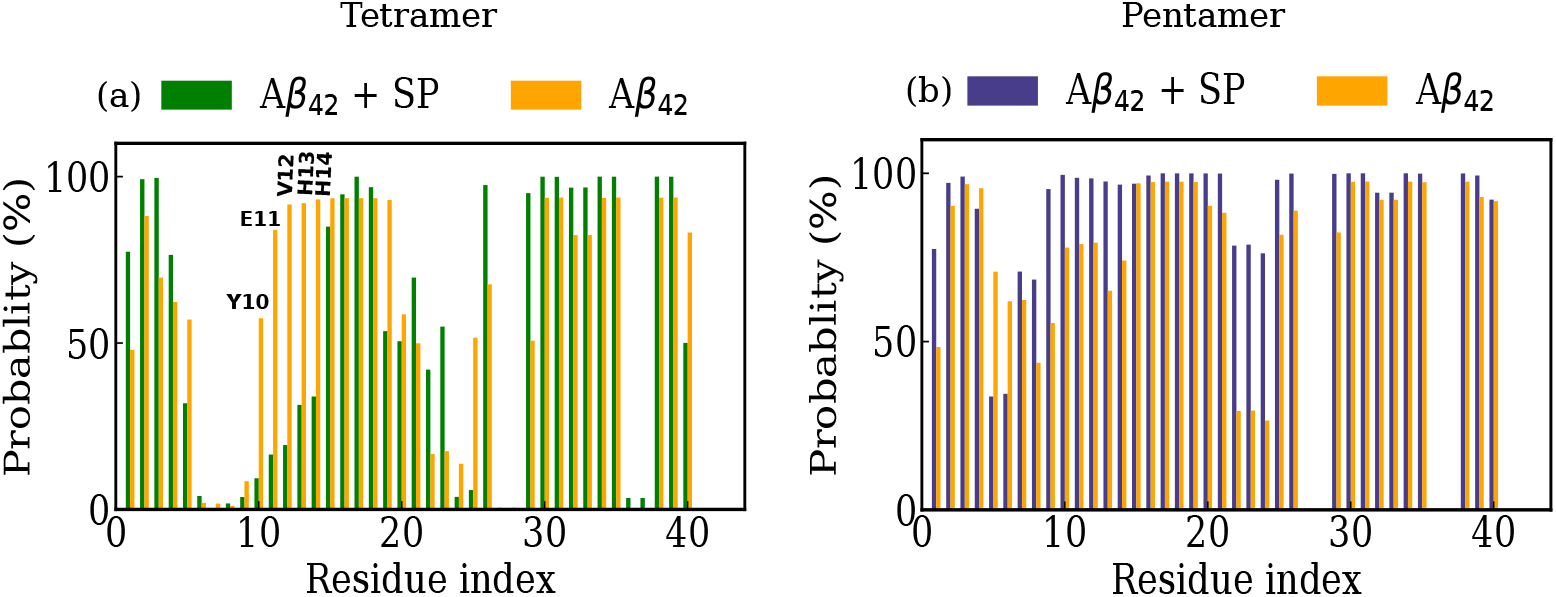
Residue-wise *β* sheet probability of A*β*_42_ protofibril for (a) Tetramer and (b) Pentamer. The *β* sheet probability calculations were performed on the last 200 ns of the trajectory.

### SP Dismantle the Salt Bridge Network in Tetramer and Pentamer, But Tetramer Shows Higher Degree of Disruption

Salt bridges plays a prominent role in stabilizing the A*β* protofibril’s structure. The presence of a salt bridge as an essential structural feature has been found across all the different morphological species of A*β* protofibril reported in the literature(*16, 35–37*). The salt bridge in A*β*_42_ protofibril(used in the present study), forms between of lysine(K28) and of alanine(A42). These salt bridges contributes to the stability of core-3 by stabilizing the C-terminus(Figure 1c and Figure 4g) of all the chains in the protofibril. To quantify the effect of SP binding on the stability of the salt bridges, we calculated the probablity density function(PDF) of the distance between the center of mass of 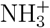 group of K28 and COO^−^ group of A42. Figure 4a and b shows PDF of intra-chain salt bridge distance for tetramer and pentamer respectively. The tetramer PDF in control system(Figure 4a) shows a sharp and high peak at around 0.45 nm with an average probability value of 0.85. The presence of a narrow and sharp peak in PDF indicate a stable network of salt bridges for tetramer in conrol system. On the other hand, PDF curve for the ligand system is spread across the higher salt bridge distances and also the area under the curve is much wider with a smaller peak(probability value = 0.2) at around 1.4 nm. These observations suggests that the intra-chain salt bridge network of tetramer is critically destabilized in the presence of SP. In case of pentamer(Figure 4b), we observed a stable salt bridge network for the isolated A*β*_42_ protofibril, indicated by two sharp peaks, one at 0.36 nm and a higher peak at 0.4 nm. A comparison between PDFs of tetramer and pentamer in the control system reveals that the intra-chain salt bridge network in pentamer is more stable than tetramer, since the dominant peak for pentamer’s PDF appears at a smaller distance(0.4 nm) than of tetramer(0.45 nm). The PDF of pentamer in ligand system shows a single distinctive peak at around 0.8 nm with a probability value of 0.6, also the area under ther curve of ligand’s PDF is broader than control’s. These findings suggests that the SP effectively disrupts the intra-chain salt bridge network in the pentamer, however, the disruption is not as severe as in tetramer. The pentamer’s PDF also explains why average RMSD for region-3 shows an increase while region-1 and region-2 shows a decrease with the SP binding(Figure 2d). Since PDF shows that SP binding disrupts salt bridge network in pentamer and both the residues(K28, A42) participating in the formation of the salt bridge lies in region-3, hence, this disturbance get reflected in the RMSD.

**Figure 4:**
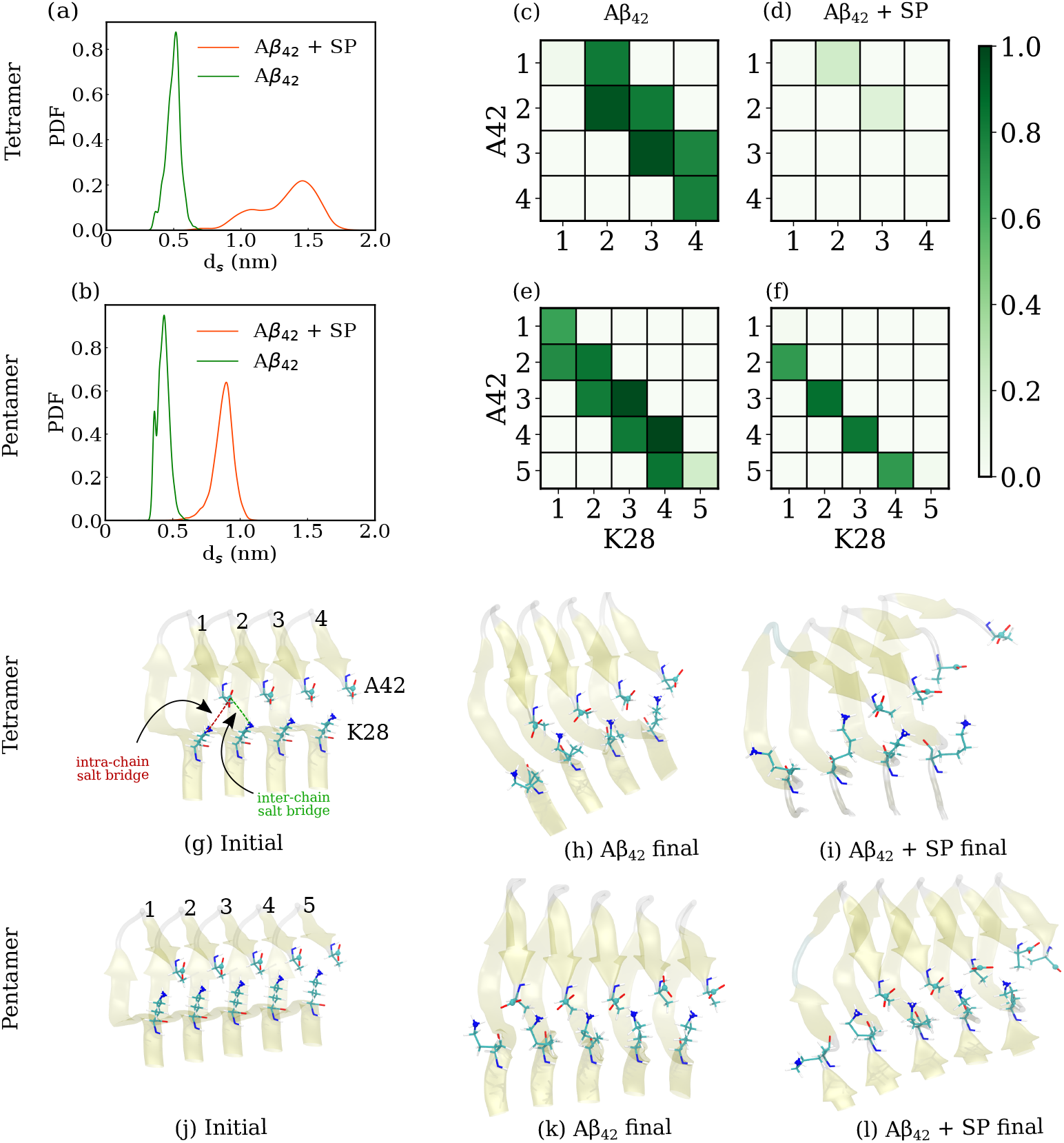
Effect of SP binding on inter-chain/intra-chain K28-A42 salt bridge. Probability distribution function of the intra-chain distance between K28 and A42 for (a) Tetramer and (b) Pentamer. Contact probability maps for all the possible inter-chain and intra-chain K28-A42 salt bridges in tetramer: (c) control system (d) ligand system and pentamer: (e) control system (f) ligand system. Snapshots of salt bridges at the start and end of simulation for (g-i) tetramer and (j-l) pentamer.

We also investigated the effect of SP binding on the stability of inter-chain salt bridges between the adjacent chains of A*β*_42_ protofibril. We calculated the probability contact maps of intra-chain and inter-chain salt bridges of tetamer and pentamer in control and ligand systems(Figure 4{*c* − *f*}). As shown in the Figure 4c, the contact map for tetramer in the control system is well populated indicating a stable network of intra and inter salt bridges. Although, the contact map suggests a weaker intra-chain salt bridge for chain-1 indicated by a very light color(almost blank) in the grid (1,1), it is comepnsated by a strong inter-chain salt bridge between A42 of chain-1 and K28 of chain-2 suggested by the strong green color of the grid (1,2). In the contact map for tetramer in the ligand system(Figure 4d) except for grid (1,2) and (2,3) showing a very light color, all grids are blank reflecting a complete disruption of both intra and inter salt bridge network of the tetramer in the presence of SP. In case of pentamer, a comparison between the contact map for control(Figure 4e) and ligand(Figure 4f) system reveals that the SP binding disrupts the salt bridge network in pentamer, however, only intra-chain bridges are effected and the inter-chain salt bridges remain intact. A visual inspection of the initial and final states of the trajectories of control and ligand systems(Figure 4g-l) corroborates well with the PDF and contact map analysis. The tetramer’s final state(Figure 4 shows a completely disordered salt bridge network, while the pentamer’s a slight disruption of the salt bridge network.

### SP Destabilizes the Hydrophobic contacts in Tetramer and Pentamer, Critically in Case of Tetramer

The three hydrophobic cores(Figure 1c) are hallmark of LS-shaped A*β*_42_ protofibril. These cores are stabilized by the contacts formed between the hydrophobic residues present in these cores. The residues involved in hydrophobic interactions in the three cores are core-1: (A2, F4, L34, V36), core-2: (L17, E19, I31), and core-3: (A30, I32, M35, V40). To examine the perturbation caused to these contacts due the presence of SP, we computed pairwise contacting frequency of these residues for each system(control and ligand). Figure 5 shows the contact maps for all the systems. As shown in the Figure 5a the tetramer in the control system have strong contacts in all three cores indicated by the high contacting frequency observed for the residues in these cores. Alternatively, the contact map for tetramer in the ligand system(Figure 5b) shows weaker contacting freqency for the key residue pairs in all three hydrophobic cores. This indicates that SP has caused disruption of key hydrophobic contacts in the tetramer of A*β*_42_ protofibril. In particular, the contacting frequency for the residue pairs F4-L34, F4-V36 in core-1, L17-I31, E19-I31 in core-2, and A30-V40, I32-V40 in core-3 has greatly decreased. Core-3 observes the most drastic disruption of the contacts among all the cores of the tetramer, all the dark colors(implying high contact fequency) in the grids of core-3 box (red box) of the control system(Figure 5a) have almost vanished in the ligand system(Figure 5b). Moreover, a comparison of contacts in core-3 between control(Figure 5a) and ligand(Figure 5) system also reveal the critical disruption of intra-chain salt bridge network(K28-A42) in tetramer, which is in agreement with the results obtained in the analysis of salt bridges in the previous section. However, the hydrophobic contacts A2-V36 and M35-V40 of the core1 and core-3 of tetramer respectively, are unaffected by the presence of SP and remains intact. The contact analysis of tetramer reveals that SP has caused significant disruption of key hydrophobic contacts in all the three cores of the protofibril. Figure 5c shows the contact map of pentamer in the control system. The contact map reveals weaker interactions between the hydrophobic residues of core-1 for the pentamer of isolated protofibril. The only residue pair showing substantial contacting frequency in the core-1 is A2-V36. Although, the contact map for pentamer in the control system revealed weaker hydrophobic interactions in core-1, the presence of SP further weakend these interactions, as can be seen in Figure 5b. The core-2 of the pentamer shows stable contacts indicated by high contacting frequency of residue pairs inside the core-2 box(green box)(Figure 5c). Although, there is a decrease in contacting frequency for many residue pair in core-2 in the presence of SP as can be seen in the Figure 5d, the key hydrophobic contacts L17-I31 and E19-I31 remains unaffected. Finally, the bold contacting frequencies of the residues in core-3 of the pentamer(red box in Figure 5c) suggests a stable core region in the control system.

**Figure 5:**
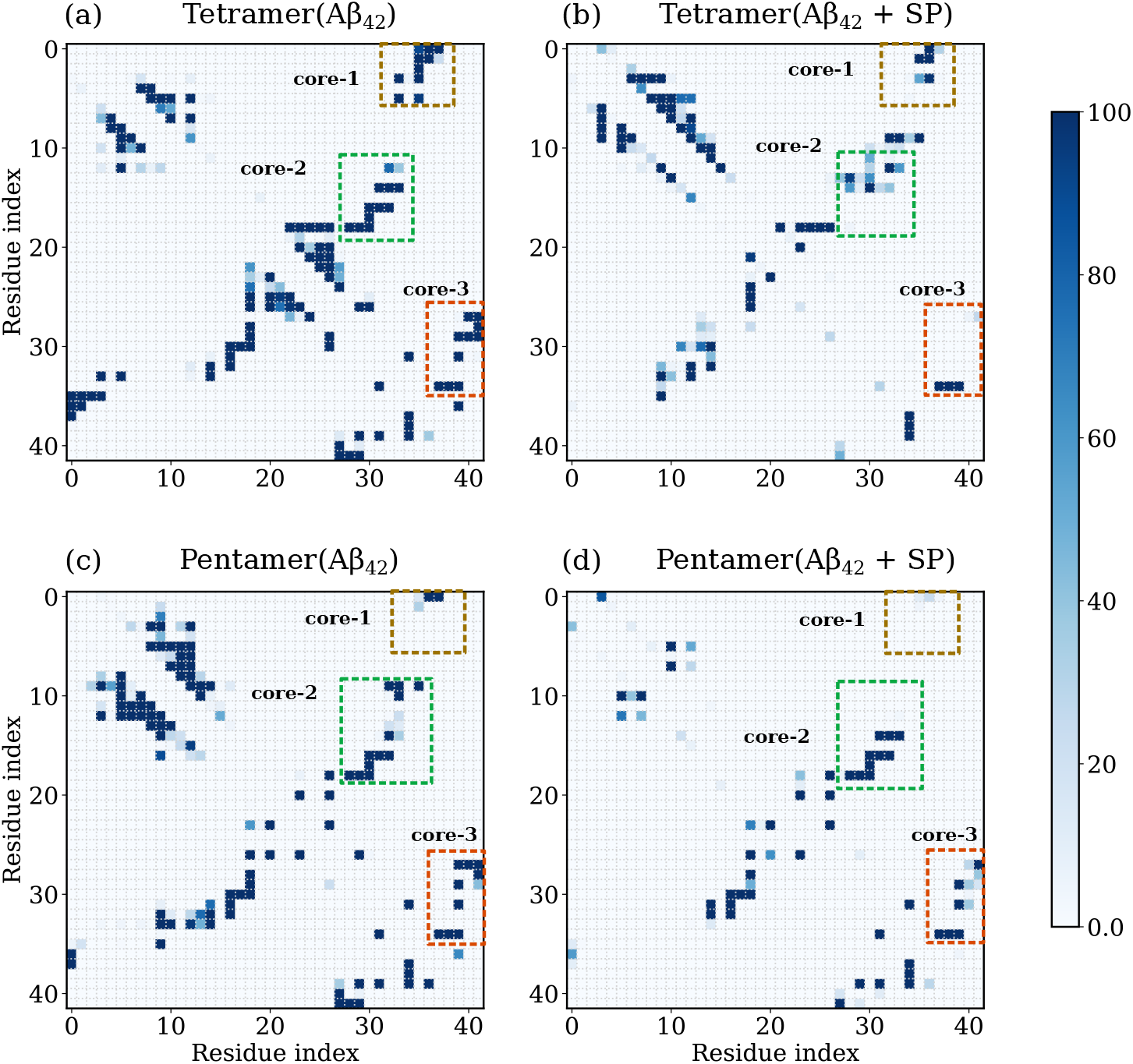
Residue-residue contacting frequency maps for tetramer in (a) control system (b) ligand system and pentamer in (c) control system and (d) ligand system. Residue contact pairs for core-1, core-2 and core-3 of A*β*_42_ protofibril are highlighted in brown, green and red dashed boxes respectively. The contacting frequency maps were computed with last 200 ns of the trajectory.

The presence of SP although influenced some residue pairs in core-3(red box in Figure 5d), however, the critical hydrophobic contacts A30-V40, I32-V40 and M35-V40 remain unaltered. Moreover, the contact maps also shows that the intra-chain salt bridge(K28-A42) of the pentamer, which is also a crucial stabilizing interaction of core-3,stay almost undisturbed by the SP binding. This observation for pentamer’s intra-chain salt bridges is in corroboration with the salt bridge analysis in the previous section.

### Binding Site Analysis and Governing Interactions

So far, we discussed the effect of SP on the structural integrity of A*β*_42_ protofibril, and the results demonstrated that the fibrillar structure is severely disrupted due to SP’s presence. However, it is also noteworthy that the pentamer’s fibril structure is relatively stable and unaffected by the SP’s presence, while the tetramer suffers a drastic loss of *β* sheet content and disruption of salt bridges, this suggests that the SP is interacting differently with pentamer and tetramer. Hence, to identify the specific binding sites of SP on tetramer and pentamer and decipher the nature of these interactions, we calculated residue-wise average interaction energies(Figure 6a and b) and contact numbers(Figure 6c and d) between A*β*_42_ protofibril residues and SP molecules.

**Figure 6:**
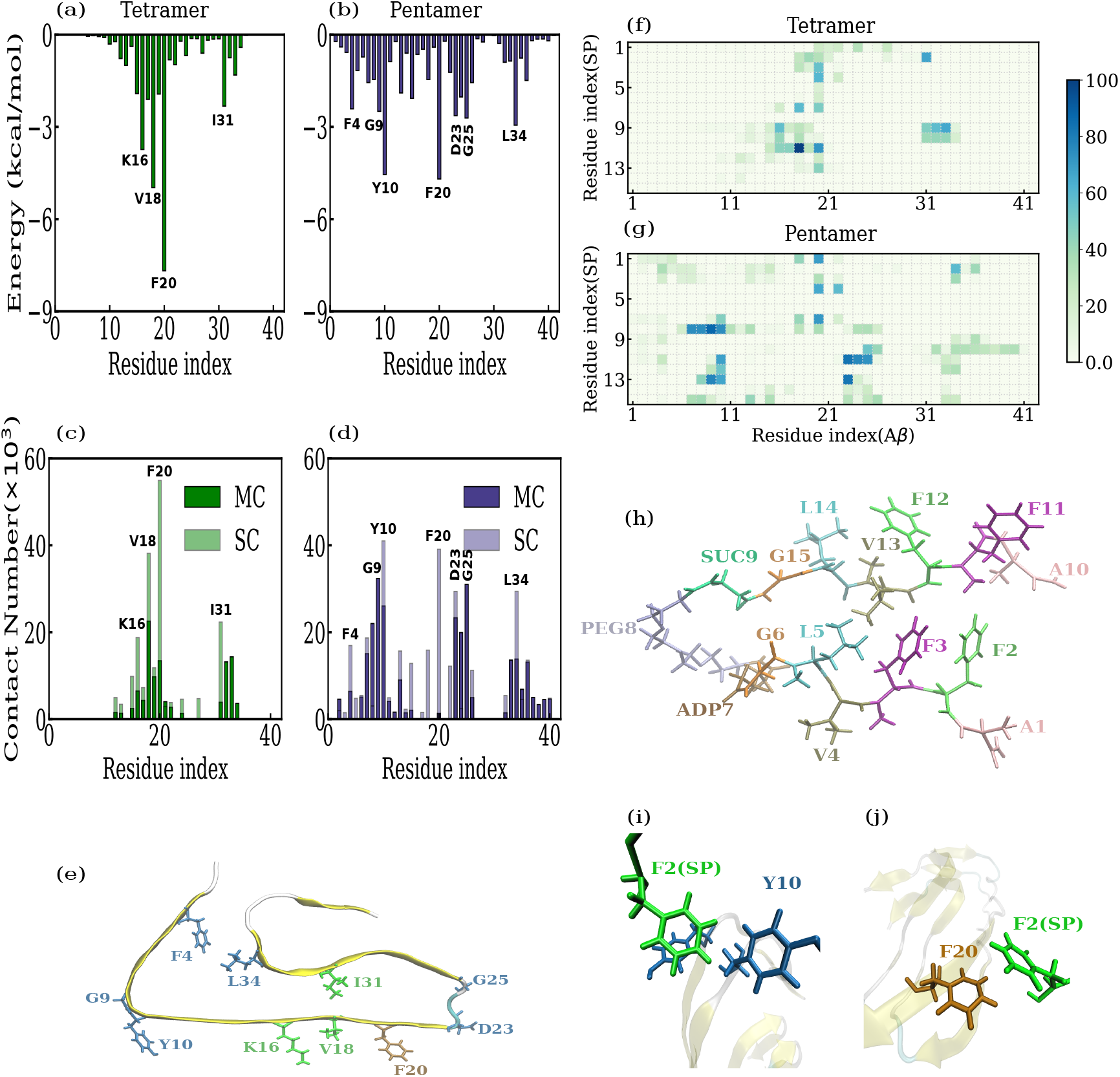
Interaction energy between each residue of (a) tetramer and (b) pentamer and SP. Number of contacts between main-chain(MC) and side-chain(SC) of each residue of (c) tetramer and (d) pentamer and SP. Visual representation of the key interacting residues of tetramer(green), pentamer(blue), and residues common to tetramer and pentamer(brown). Contacting frequency maps between residue pair of SP and (f) tetramer and (g) pentamer. (h) Licroice representation of the structure of SP. The residues are shown in different colors and are annotated with residue name and number. Snapshot of *π* − *π* interaction between one of the phenylalanine(F2) of SP with (i) Y10 of pentamer and (j) F20 of tetramer.

We can note from Figure 6a that the tetramer’s residues showing high binding affinity with SP are F20, V18, K16, and I31 in the order of decreasing interaction energy. The presence of phenylalanine, valine, and isoleucine in the above list of the key interacting residues reveals the prominent role of hydrophobic and aromatic residues in the SP binding. Interestingly, the residues K16, V18, and F20 belong to the amyloidogenic core region(KLVFFA), and the subunit LVFFA from this region is the primary ingredient in the design of SP. The high binding affinities of these residues indicate that the SP can recognize the subunit in A*β*_42_ protofibril and bind strongly to it. Figure 6c shows the number of contacts each residue of tetramer has with SP. The contact number analysis tells a similar story as the interaction energies for tetramer. The residues with high contact numbers are the same with high interaction energies and in the same order. Additionally, the contact analysis highlights the role of side-chain interaction in the binding of SP. As can be observed from Figure 6c, all the key interacting residues have higher side chain contacts, indicating the presence of aromatic and hydrophobic interactions with SP. For pentamer, the key interacting residues involved in SP binding, as revealed by interaction energy(Figure 6b and contact(Figure 6d) analysis are: F20, Y10, L34, D23, G25, G9 and F4 in the order of decreasing interaction energy and contact number. Again, the residues in pentamer that show high affinity towards SP are primarily aromatic(F20, Y10, and F4) and hydrophobic(L34). Also, the contact analysis for pentamer(Figure 6d) shows that the key interacting residues exhibit higher side-chain contacts.

Although the number of key interacting residues in the case of pentamer is higher than tetramer, a comparison of interaction energies(Figure 6 a and d) and number of contacts(Figure 6 c and d) between tetramer and pentamer shows that tetramer exhibits higher binding affinity towards SP, particularly F20 of tetramer displays interaction energy and contact number that are at least twice than any residue in pentamer. Moreover, as explained above, the tetramer residues with high binding affinity to SP belong to the amyloidogenic core region, while these residues in pentamer show relatively weaker interaction with SP. Furthermore, the markedly strong interaction of F20 with SP suggests weakening of the *π* − *π* interaction between the benzene rings of F20 residue of adjacent chains of the tetramer. And as these interactions play a central role in the formation of A*β* fibrils(*38, 39*), any perturbation to these interactions will cause modification of the fibril’s structure. Thus, all the above observations explain the pronounced destabilization of fibrillar structure in the case of tetramer while not so much for pentamer. Figure 6e shows the relative position and orientation of the key interacting residues in the LS-shaped A*β*_42_ protofibril. We can observe from Figure 6e that residues with high binding affinity to SP have the side chains oriented outside the surface of the protofibril. The preferential binding of SP to the residues with side chains oriented outwards can be attributed to the complex hairpin-like chemical structure of the SP(Figure 6h). Due to the structural restrictions, it would be difficult for SP to interact with the residues with side chains facing inward; therefore, it preferentially binds to the outward-facing residues. However, Figure 6e shows few inward-facing residues(F4, I31, and L34) also, showing favorable interactions with SP. These residues must belong to the corner chains of the protofibril(tetramer/pentamer), where they are easily accessible to SP for interactions. Still, these residues exhibit relatively weaker interactions with SP than the outward-facing residues, as observed from the interaction energy and contact number plots.

Residue-wise interaction energy and contact analysis provided information about the residues of A*β*_42_ protofibril participating in the binding with SP. However, to gain a deeper understanding of SP-A*β* binding, we computed contacting frequency maps(Figure 6f and g) residue pairs between A*β*_42_ and SP. Figure 6h shows different residues of SP in different colors and their corresponding name and number. Figure 6f shows a contact map between the residues of SP and tetramer. The residue K16 of tetramer shows a high contacting frequency with SUC9(succinyl), alanine(A10), and phenylalanine(F11) residues of SP. A strong binding between K16(positively charged side-chain) of A*β* and F11(aromatic side-chain) of SP suggest the presence of cation-*π* interaction. L17(leucine) shows a high contact frequency with only F11 of SP, indicating a hydrophobic interaction between them. V18(valine) of tetramer shows pronounced contacts with F11, ADP7(adipoyl), F2, and F3 of SP. All these residues are hydrophobic/aromatic, implying a strong hydrophobic interaction with V18. F19 of tetramer exhibits high contacting frequency with F2, and a medium one with SUC9 and F11 suggests the presence of *π* − *π* and hydrophobic interactions. As evident from interaction energy and contact analysis for tetramer, F20 shows the highest number of high-frequency contact pairs with SP residues. The residues of SP in contact with F20 are F2, F3, V4, ADP7, F11, V13, and F12. As can be observed, many aromatic and hydrophobic residues show close interaction with F20, pointing towards the presence of *π* − *π* and hydrophobic interactions between F20 and SP. The residue I31 exhibits aromatic and hydrophobic contacts with F2, SUC9, and A10. Finally, I32 and G33 show marked contacts with SUC9 and A10. Figure 6j shows a snapshot depicting *π* − *π* interaction between F20 of tetramer and F2 of SP.

A superficial overview of the contact map for pentamer and SP(Figure 6g) shows that pentamer has more binding sites than tetramer, evident from interaction energy and contact number analysis. F4 of pentamer shows strong interaction with F2, F3, and F11, implying *π* − *π* interaction with each of these residues with F4. Residues D7(aspartate) and S8(serine) show strong interaction with PEG8(polyethylene glycol) of SP and weak interaction with A1(alanine). These contacts are mainly with the main chain of the pentamer’s residues, as observed from the contact number analysis(Figure 6d), and hence are primarily hydrophobic. Similarly, G9 of pentamer also shows strong main-chain contacts with PEG8 and V13(valine) of SP. Further, Y10(tyrosine) of pentamer exhibits pronounced contacts with F11, F12, V13, and PEG8, indicating strong hydrophobic and *π* − *π* interaction; Y10 also shows a mild interaction with ADP7. For the V18 residue of pentamer, we observe two contacts with A1 and ADP7 with medium contact frequency and a weak one with PEG8. The F20 of pentamer establishes strong interactions with A1, V4, and ADP7 of SP while interactions of medium strength with F3 and PEG8. The F20 has strong hydrophobic but a weaker aromatic interaction with SP. Residues D23, V24, and G25 have high contact frequency with F11 of SP, and D23 also shows firm contact with V13. Finally, L34 of pentamer shows strong hydrophobic contacts with F2 and considerable contacts with F3.

Overall, the residue-wise interaction energy and contact number analysis revealed crucial residues in tetramer/pentamer participating in binding with SP. Although pentamer showed higher binding residues, tetramer’s residues exhibited stronger interaction, particularly in the amyloidogenic core region. The contact maps between the residues of SP and tetramer/pentamer identified the residue pairs in close and sustained contact, consequently revealing the nature of the interaction between SP and A*β*_42_ protofibril. We found a range of interactions at play; however, the *π* − *π* and hydrophobic interaction were the most dominant mode of interaction.

## 3 Conclusions

In this study, we investigated the effects and molecular mechanism of a hairpin-like synthetic paratope binding on a preformed LS-shaped A*β*_42_ protofibril using MD simulations. Our simulations show that SP can destabilize the overall structural integrity of A*β*_42_ protofibril. The RMSD analysis revealed that the tetramer of the A*β*_42_ protofibril is severely disrupted, while pentamer remains almost unaffected by the presence of SP. The region Y10-N27 in the tetramer suffered the maximum structural distortion, while D1-G9 and K28-A42 also displayed significant deviation from their native structures in the presence of SP. The SP binding caused a considerable loss of *β* sheet content in tetramer, particularly for the segment Y10-H14, while pentamer showed no serious changes in its *β* sheet due to the SP binding. The K28-A42 salt bridge is a crucial factor in the A*β*_42_ protofibril’s structural stability, and our simulations showed that SP causes severe damage to intra-chain and inter-chain salt bridges in the case of the tetramer. The contacts in the three hydrophobic cores of A*β*_42_ protofibril are strongly affected by SP in case of tetramer, especially the key hydrophobic contacts in these cores such as core-1:[(F4-L34), (F4-V36)], core-2:[(L17-I31), (E19-I31)], and core-3:[(A30-V40), (I32-V40)] are heavily disrupted due to SP binding.

The binding site analysis revealed interesting details about the differences in the binding of SP with tetramer and pentamer and the binding mechanism of SP with A*β*_42_ protofibril. Interestingly, for tetramer, the SP was able to self-recognize the LVFFA region on the protofibril, which is the key component of the SP’s design, and the SP binds strongly to this region with F20, V18, and K16 as the participating residues. The key binding residues for pentamer are F20, Y10, D23, L34, and F4. Although pentamer shows more binding sites than tetramer, the pentamer’s residues have much weaker binding with SP, which explains why we do not observe any significant disaggregation of fibrils in the case of pentamer. The SP preferentially binds to the residues with side chains oriented outward from the LS-shaped A*β*_42_ protofibril surface. The dominant modes of interaction identified between A*β*_42_ protofibril and SP are *π* − *π* and hydrophobic interaction. Our study provides interesting atomistic details about SP binding onto A*β*_42_ protofibril. It also sheds light on the disaggregation mechanism of the fibrils due to SP binding. The results of our study can prove to be useful in developing therapeutics for AD.

## 4 Methods

### Control System Design

The initial structure of the A*β*_42_ protofibril was obtained from the protein data bank entry: 5OQV, which is derived using cryo-EM(*16*). The chosen model contains nine chains of LS-shaped conformation of full-length A*β*_1− 42_ monomers. The nine chains are two multimers: tetramer and pentamer(Figure 1a). The nonamer(nine chains) protofibril is solvated with water molecules in a cubic box with an edge length of 9.4 nm. The box length is designed such that the closest distance between the protofibril and box boundary is at least 1.2 nm. The solvated system is further neutralized by adding twenty-seven sodium(Na^+^) ions. The final system contains 78120 atoms in total.

### Ligand and Ligand System Design

The structure of the ligand(SP)(Figure 1d) was obtained from the work by Paul et al.(*34*). As shown in Figure 1d, the structure of SP has two copies of short peptide fragment(LVFFA) from the amyloidogenic core region of A*β*, connected with a flexible turn region made with adipoyl, polyethylene glycol, and succinyl, to create a *β* hairpin-like structure. For a detailed account of the design and synthesis of SP, the reader is referred to the work by Paul et al.(*34*). The atomistic model of SP for our MD simulations was constructed, optimized, and energy minimized using Avogadro(*40*). The CGenFF program(*41*) was used to generate CHARMM-compatible forcefield parameters and partial charges for the SP molecule. These parameters were further refined using the forcefield toolkit(*42*) in the visual molecular dynamics(VMD) program(*43*). Subsequently, a merged system is constructed with SP molecules placed randomly around A*β*_42_ nonamer at a distance of 1.2 nm from the non-amer and from each other(Figure 1b). The number of SP molecules in the ligand system was kept in a ratio of 1:1 to the number of A*β*_42_ chains. The merged system is solvated with water molecules in a rectangular box of dimension 9.2 × 10.5 × 12.7 nm^3^. The box dimensions were so designed that any atom of SP or A*β*_42_ protofibril is at least 1.2 nm away from the simulation box’s boundary. Further, 27 sodium ions were added to the system to maintain charge neutrality. The final ligand system contains 114876 atoms.

### Simulation Details

All simulations were performed using the NAMD 2.14 MD simulation package(*44*). The Charmm36 force field(*45*) was used for the A*β* fibril and TIP3P water model(*46*). All systems were energy minimized using 5000 steepest descent steps. The systems were then equilibrated for 100 ps using the canonical (NVT) ensemble, followed by a further 100 ps equilibration simulation with the isobaric-isothermic (NPT) ensemble. The production runs for all the systems were performed in the NPT ensemble. All the covalent bonds involving hydrogen atoms in water molecules were constrained using SETTLE algorithm(*47*), and those except water molecules were constrained with SHAKE algorithm(*48*). We used a time step of 2 fs for integrating the equations of motion. Particle mesh Ewald (PME) method(*49*) was used to calculate long-range electrostatic interactions with a real space cutoff of 1.2 nm. The van der Waals (vdW) interactions were calculated using a cutoff of 1.2 nm. All MD simulations were performed at a temperature of 310 K, and the temperature control was achieved using a Langevin thermostat(*50*). A constant pressure of 1.01325 bar was maintained in an NPT ensemble by using Nosé-Hoover Langevin barostat(*51, 52*).

### Analysis Methods

All the analysis was performed using MDTraj(*53*), VMD(*43*), NAMD(*44*), and our in-house developed Tcl, python, and bash codes. The RMSD of C*α* atoms of A*β*_42_ protofibril was computed with MDTraj. The initial structure of A*β*_42_ protofibril was taken as a reference, and the RMSD for each chain in the tetramer/pentamer was calculated and averaged over all the chains in the tetramer/pentamer. The secondary structure calculations were performed with the STRIDE algorithm(*54*) in VMD. The K28-A42 salt bridge plays a critical role in the structural stability of A*β*_42_ protofibril. The K28-A42 salt bridge is said to exist if the distance between positively charged Nζ of K28 and negatively charged C*γ*-carboxylate of A42 is within 0.4 nm. Residue-wise interaction energies were extracted using the *pari-interaction* facility in NAMD. All the interaction energies reported here are vdW energies unless otherwise stated. In the analysis of contacts between the residues of tetramer/pentamer(contact maps in Figure 5), the contacting frequency is computed between two residues separated by two peptide bonds. The contacting frequency was calculated for residue pairs in a single chain and then averaged over all the chains in the tetramer/pentamer. A contact is said to exist if the distance between two non-hydrogen atoms is within 0.5 nm. All the visualizations were performed with VMD(*43*).

## Acknowledgement

Authors thanks Indian Institue of Technology Guwahati for providing High-Performance computing facility “Param-Ishan” for performing the simulations.

## Notes

### Competing Interest Statement

The authors have declared no competing interest.

